# The time-course of component processes of selective attention

**DOI:** 10.1101/511022

**Authors:** Tanya Wen, John Duncan, Daniel J Mitchell

**Affiliations:** Medical Research Council, Cognition and Brain Sciences Unit, University of Cambridge, 15 Chaucer Road, Cambridge, CB2 7EF, United Kingdom; Department of Psychology, University of Oxford, Oxford, UK

**Keywords:** selective attention, MEG/EEG, decoding, attentional template, visual processing

## Abstract

Attentional selection shapes human perception, enhancing relevant information, according to behavioral goals. While many studies have investigated individual neural signatures of attention, here we used multivariate decoding of electrophysiological brain responses (MEG/EEG) to track and compare multiple component processes of selective attention. Auditory cues instructed participants to select a particular visual target, embedded within a subsequent stream of displays. Combining single and multi-item displays with different types of distractors allowed multiple aspects of information content to be decoded, distinguishing distinct components of attention, as the selection process evolved. Although the task required comparison of items to an attentional “template” held in memory, signals consistent with such a template were largely undetectable throughout the preparatory period but re-emerged after presentation of a non-target choice display. Choice displays evoked strong neural representation of multiple target features, evolving over different timescales. We quantified five distinct processing operations with different time-courses. First, visual properties of the stimulus were strongly represented. Second, the candidate target was rapidly identified and localized in multi-item displays, providing the earliest evidence of modulation by behavioral relevance. Third, the identity of the target continued to be enhanced, relative to distractors. Fourth, only later was the behavioral significance of the target explicitly represented in single-item displays. Finally, if the target was not identified and search was to be resumed, then an attentional template was weakly reactivated. The observation that an item’s behavioral relevance directs attention in multi-item displays prior to explicit representation of target/non-target status in single-item displays is consistent with two-stage models of attention.

## 1. Introduction

Our perception of the world is constantly shaped by attentional selection, enhancing relevant over irrelevant information, to achieve our behavioral goals. Effective selection begins from a flexible description, often called the attentional template, of the object currently required (Duncan and Humphreys, 1989; Bundesen, 1990). Much evidence suggests that attentional selection is then achieved through a process of biased, integrated competition across a broad sensorimotor network (Duncan et al., 1997). As objects in the visual input compete to dominate neural activity, the degree to which they match the attentional template determines their competitive advantage (Desimone and Duncan, 1995; Beck and Kastner, 2009).

Attention is often characterized as an emergent property of numerous neural mechanisms (Desimone & Duncan, 1995; Hopf et al. 2005), with different mechanisms dominating as successive stages of selection (Eimer, 2015). Therefore, while many studies have investigated the time-course of individual neural signatures of attention in humans and animal models, it is informative to compare multiple components of the selection process within the same paradigm. Recently, there has been much interest in the use of MEG/EEG for real-time decoding of cognitive representations in the human brain (Stokes et al., 2015). Here, we used simultaneous MEG/EEG to examine the time-course and content of different components of attentional selection. We combined single-item and multi-item search displays with different types of distractors to allow multiple aspects of information content to be decoded from the neural signal, distinguishing distinct components of attention as the selection process evolved.

The behavioral relevance of stimuli was manipulated by starting each trial with one of two auditory cues, indicating the relevant visual target object on this trial. Participants were then presented with a series of visual displays of 4 possible types: a 1-item display of the target (T), an inconsistent non-target (Ni; which was associated with the other cue and served as a target for other trials), a consistent non-target (Nc; which was never a target), or a 3-item display with all items presented simultaneously (see Figure 1 for an illustration). The use of inconsistent non-targets allowed representation of target status to be distinguished from representation of stimulus identity. The inclusion of 3-item displays allowed competitive representation of target location and target identity to be quantified under matched visual input. The use of consistent non-targets amongst a stream of choice displays allowed decoding of attentional template reactivation in preparation for a subsequent display. Participants made a button press whenever they detected a rare brightening of the target item. Requiring responses only for conjunctions of identity and brightening allowed response trials to be excluded from the analysis and attentional selection assessed on trials without an overt response. Using multivariate decoding analyses, we asked which component processes of attentional selection are visible in the MEG/EEG signal over time.

**Figure 1.**
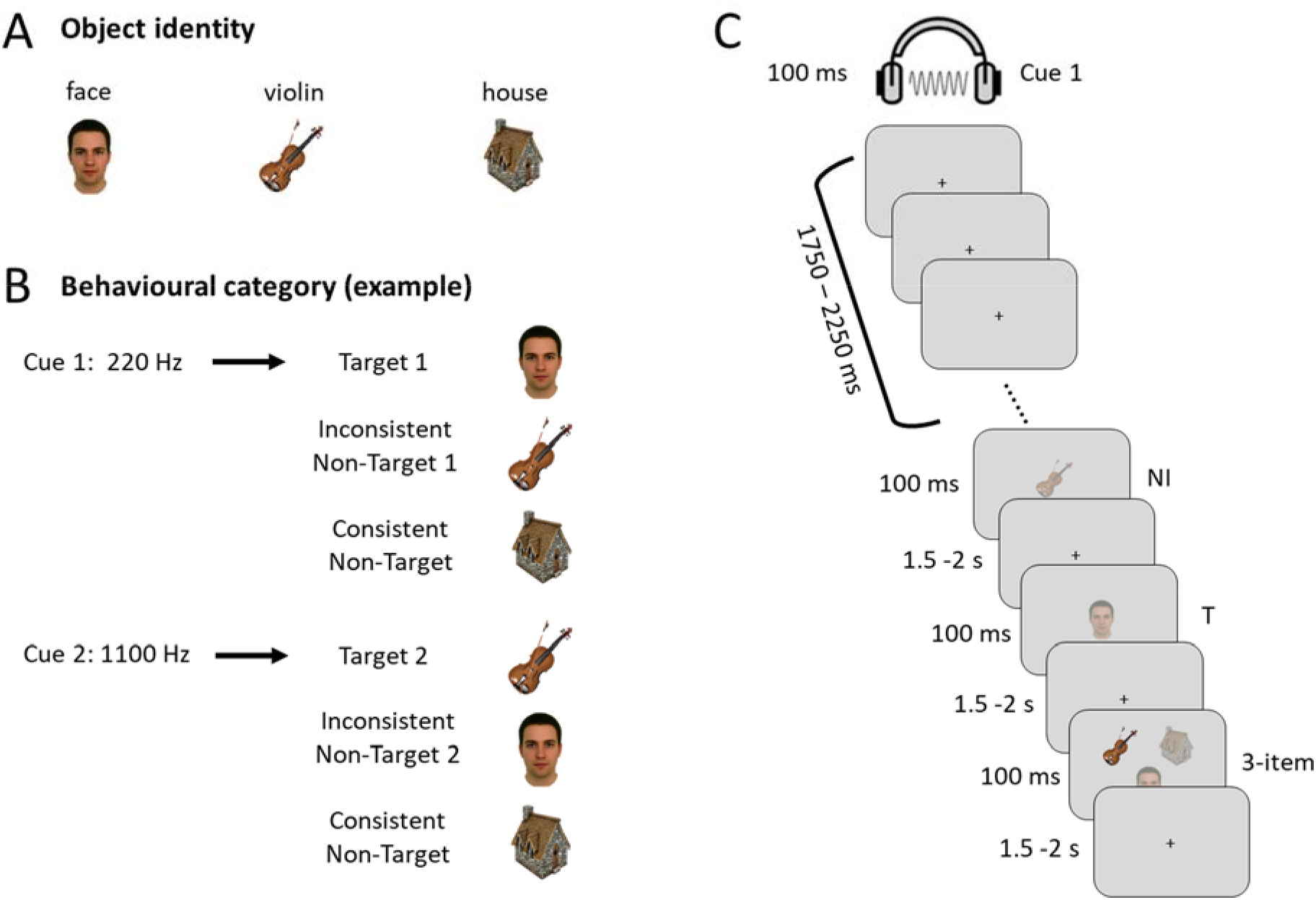
Stimuli and experimental paradigm. (A) The 3 objects used in the experiment. (B) An example of how the two auditory tones could be paired with the three objects. This results in two items that serve as targets (T) for one cue, and non-targets (Ni) for the other cue, and the third item serving as a consistent non-target (Nc). The pairings between the tones and the objects were counterbalanced across participants. (C) An example trial illustrating the experimental paradigm. At the beginning of each trial, an auditory cue indicated the target for that trial. After a delay, this was followed by three visual displays. Participants were asked to make an immediate button press if a brightening of the target stimulus was detected.

First, we examined representation of the attentional template. One possibility is that, when a cue indicates the relevant target object, some sustained signal will be set up in neurons selectively responsive to that object (Chelazzi et al., 1993; Puri et al., 2009; Kok et al., 2013). fMRI decoding studies have shown cross-generalization between attentional templates and sensory responses to the corresponding objects (e.g., Stokes et al., 2009; Peelen and Kastner, 2011), supporting a tonic activation of visual representations. However, corresponding results tend to be weak or non-existent in electrophysiological recordings (Stokes et al., 2013; Myers et al., 2015; Wolff et al., 2015), and where they have been found, they may appear only very briefly prior to the target stimulus (Myers et al., 2015; Kok et al., 2017). Indirect measures of attentional templates, derived from ERP components, demonstrate that search templates are not continuously active but are transiently activated in preparation for each new search episode (Grubert and Eimer, 2018). Recently, it has been proposed that template storage may sometimes be “silent”, perhaps encoded in changed synaptic weights rather than sustained firing (Stokes, 2015). To examine template coding, holding visual input constant, we analyzed data from the period between cue and displays, and during subsequent presentation of Nc stimuli.

Second, we were interested in the process of target selection itself. Comparing target and non-target stimuli shows strong differences both behaviorally and neurally (Duncan, 1980; Hebart et al., 2018). Attending to a relevant visual object produces strong, sustained activity across many brain regions (Desimone and Duncan, 1995; Sergent et al., 2005; Dehaene and Changeux, 2011), reflecting encoding of its multiple visual properties and implications for behavior (Wutz et al., 2018). In the presence of multiple stimuli, neural responses are initially divided amongst the competing sensory inputs and later become replaced by a wide-spread processing of the behaviorally critical target (Duncan et al., 1997; Kadohisa et al., 2013). On 1-item trials, we focused on the response to the T and Ni stimuli, to quantify the representation of object identity (e.g., face vs. house) regardless of status as target or non-target, as well as representation of behavioral category (T vs. Ni) regardless of object identity. On 3-item trials, we quantified the encoding of target location and target identity, to assess preferential processing of target features when multiple items compete for representation.

## 2. Methods

### 2.1 Participants

Eighteen participants (9 males, 9 females; age range: 18-30 years, mean = 24.4, SD = 3.8) took part in the study, recruited from the volunteer panel of the MRC Cognition and Brain Sciences Unit. Two additional participants were excluded from the analysis due to technical problems (one could not do the MRI; another was excluded due to an error in digitizing the EEG electrodes). EEG data for 4 participants were excluded from the MVPA analysis due to a technical issue (a test signal used during hardware checkup was not removed). All participants were neurologically healthy, right-handed, with normal hearing and normal or corrected-to-normal vision. Procedures were carried out in accordance with ethical approval obtained from the Cambridge Psychology Research Ethics Committee, and participants provided written, informed consent prior to the experiment.

### 2.2 Stimuli and Procedures

Participants performed two localizer tasks (auditory and visual) and an attention task (see Figure 1 for an illustration). Stimulus presentation was controlled using the Psychophysics Toolbox (Brainard, 1997) in Matlab 2014a (Mathworks, Natick, WA). Auditory stimuli were delivered through in-ear headphones compatible with the MEG recording. Visual stimuli were back-projected onto a screen placed in front of the participant, approximately 129 cm from the participant’s eyes. Each stimulus image was approximately 20 cm wide (approximate visual angle 8.8°) on a gray background. Before the start of each task, participants were given training to familiarize them with the stimuli and task rules. If a false alarm was made during any of the trials during the recording, that trial was repeated at the end of the run.

#### 2.2.1 Pattern Localizer Tasks

Auditory Localizer Task: This task was used to characterize multivariate activity patterns for high and low pitch tones used in the attention task. Participants heard a stream of intermixed high (1100 Hz) and low (220 Hz) pitch tones. On rare occasions (9% of the time), a frequency modulation would occur (modulator frequency = 20 Hz; modulation index = 2.5), and participants were instructed to press a button whenever they detected a distortion in a tone. There were 100 occurrences of each unmodulated tone and 10 occurrences of each modulated tone. The duration of each tone was 100 ms, with the beginning and ending 10 ms ramped. The inter-stimulus interval was jittered between 1000-1500 ms.

Visual Localizer Task: Similar to the auditory localizer task, this task was used to establish multivariate activity patterns for three visual stimuli (a face, a house, and a violin) used in the attention task. Participants were shown a stream of these images presented sequentially in the center of the screen for 100 ms each, with an inter-stimulus interval jittered between 1500-2000 ms. Most image displays were semi-transparent (60% opaque) on a grey background; participants were asked to make a button press whenever they detected a brighter and higher contrast version of the image (100% opaque). There were 100 occurrences of each translucent image and 10 occurrences of each brightened image.

#### 2.2.2 Attention Task

Figure 1 illustrates the stimuli used in the task, as well as the task structure. Before the start of this task, participants were trained to associate the two auditory tones with two of the three visual stimuli (the same used in the localizer tasks). This pairing resulted in the visual stimuli being categorized by behavioral relevance as targets (T: the visual stimulus paired with the current cue), inconsistent non-targets (Ni: the visual stimulus paired with the alternative cue), and consistent non-targets (Nc: never targets). All six possible mappings of two cues to three objects were counterbalanced across participants. The task was executed in runs of 90 trials. Each trial began with an auditory cue (for 100 ms), followed by a 1750 – 2250 ms fixation cross during which participants were instructed to prepare to attend for the target stimulus. Then a stream of three visual displays appeared one by one for 100 ms each, separated by 1500-2000 ms inter-stimulus intervals. Each display could be a 1-item display or a 3-item display with equal probability (order pseudorandomized, with the constraint that a 1-item display could not follow a 1-item display of the same type to minimize sensory adaptation effects). On 1-item displays, the stimulus was centered at fixation; 3-item displays contained all three visual stimuli, with the center of each stimulus 10° visual angle from fixation, arranged in an equilateral triangle with one above left, one above right, and one below. In 18 out of the 90 trials in each run, a single brightened stimulus, target or non-target, occurred pseudo-randomly in one of the 3 displays, with equal likelihood of appearing in each. For each cue type, brightenings affected T, Ni and Nc items once each on single item-trials, and twice each on 3-item displays, allowing one brightening for each of the six possible 3-item stimulus configurations. Participants were asked to attend to targets, pressing a button if they detected a brightened target (they could respond any time before the next stimulus), with no response for all other displays. Events with a brightened stimulus and/or button presses were later removed in the analysis, such that the results were not influenced by these events. The trial terminated if a button press was made, and participants were informed whether the response was a correct detection or a false alarm. A new trial began when the participant indicated with a button press that they were ready to continue. Otherwise, each of the 90 trials in each run had a full sequence of 3 displays. At the end of each run, feedback informed participants of their accuracy through the run. To discourage false alarms and equalize the number of non-response trials across conditions, trials that contained a false alarm were repeated at the end of the run. The task was repeated over 5 runs (2 participants only completed 4 runs due to time constraints).

### 2.3 Data acquisition

#### 2.3.1 Electroencephalography (EEG)

EEG data were collected from 70 Ag/AgCl electrodes mounted on an electrode cap (Easycap, Falk Minow Services, Herrsching-Breitbrunn, Germany) distributed according to the extended 10/20 system. Electrode impedances were kept below 5 kΩ. An electrode placed on the nose served as online reference while the ground electrode was placed on the right cheek. Vertical and horizontal eye movements were monitored using the electrooculograms (EOG) recorded using bipolar electrodes placed above and below the left eye and at the outer canthi of the eyes, respectively. Electrocardiography (ECG) was recorded using bipolar electrodes placed below the right collarbone and below the left ribcage. EEG data were sampled at 1000 Hz with a band-pass filter of 0.1–333 Hz. EEG and MEG data were acquired simultaneously.

#### 2.3.2 Magnetoencephalography (MEG)

MEG data were acquired using a 306 channel (204 planar gradiometers and 102 magnetometers) Neuromag Vectorview system (Elekta AB, Stockholm) in a sound-attenuated and magnetically shielded room. Data were sampled at 1000 Hz with an online band-pass filter of 0.03–333 Hz. Five Head Position Indicator (HPI) coils were attached firmly to the EEG cap to track the head movements of the participant. The locations of the HPI coils as well as the EEG electrodes were recorded with a Polhemus 3D digitizer. We also measured three anatomical landmark points (nasion, left and right preauricular points) and additional points on the head to indicate head shape and enable matching to each individual’s structural MRI scan.

#### 2.3.3 Structural MRIs

High-resolution anatomical T1-weighted images were acquired for each participant (either after the MEG session or at least three days prior to the MEG session) in a 3T Siemens Tim Trio scanner, using a 3D MPRAGE sequence (192 axial slices, TR = 2250 ms, TI = 900 ms, TE = 2.99 ms, flip angle = 9°, field of view = 256 mm × 240 mm × 160 mm, 1 mm isotropic resolution). The coordinates of the nasion, left and right preauricular points in native space were hand-marked by the experimenter, and used in the coregistration of the EEG/MEG and MRI.

### 2.4 EEG and MEG data preprocessing

The raw data were visually inspected during recording for any bad channels, which were removed (EEG: 0 - 5 across subjects; MEG: 1 - 5 across subjects). The MEG data were de-noised using Maxfilter 2.2 (Elekta Neuromag, Helsinki), with the spherical harmonic model centered on a sphere fit to the digitized head points; default settings were used for the number of basis functions and the spatiotemporal extension (Taulu & Simola, 2006). Maxfilter detected additional bad channels using the first and last 900 data samples (default threshold), and signal from all bad channels was removed and interpolated. Continuous movement compensation was applied at the same time.

Subsequent preprocessing used SPM12 (http://www.fil.ion.ucl.ac.uk/spm) and Matlab 2015a (The Mathworks Inc). Separately for EEG electrodes, magnetometers and gradiometers, independent component analysis (ICA), implemented using EEGLAB (Delorme & Makeig, 2004), was used to detect and remove components whose time-course correlated with EOG or ECG reference time-courses, and whose topography matched reference topographies associated with ocular or cardiac artefacts estimated from independent data acquired on the same system. ICA used the default infomax algorithm, with dimension reduction to 60 principal components. An independent component was removed if (1) it had the maximum absolute correlation with both a temporal and spatial reference, (2) these correlations were significant at p < 0.05, (3) the z-scored absolute correlations exceeded 2 for the spatial component, and 3 for the temporal component, and (4) it explained > 1.7% of total variance. For assessing temporal correlations only, ICA and reference time-courses were band-pass filtered between 0.1 - 25 Hz, and correlations were also repeated 1000 times with phase randomization of the reference time-course to ensure that the true maximum absolute correlation of eliminated components was greater than the 95^th^ percentile of the null distribution. EEG data were then re-referenced to the average reference.

Data were band-pass filtered between 0.1 Hz and 40 Hz (zero-phase forward and reverse 5^th^ order Butterworth filters with half-power cutoff frequencies). We note that although filtering enhances the signal-to-noise ratio of neural signals, it also spreads signal in time, distorting estimates of onset latencies. In this paper we focus on peak latencies, which are less sensitive to filtering (Luck, 2014; Grootswagers et al., 2016; van Driel et al., 2019). Data were epoched around the events of interest, time-locked to stimulus onset (from −100 ms to 1000 ms in the auditory localizer task; from −100 ms to 1500 ms in the visual localizer task; from - 100 ms to 1750 ms for the cue and delay period of the main task, and −100 ms to 1500 ms for each of the visual stimulus presentations in the main task). Time points −100 ms to 0 ms served as baseline for baseline correction – the mean signal across this window was subtracted from each time point, per epoch. Epochs that contained flat segments or high threshold artifacts (peak-to-peak amplitude greater than 4000 fT for magnetometers, 400 fT/m for gradiometers, 120 μV for EEG, or 750 μV for EOG) were marked as bad trials and were rejected. In both localizer and attention tasks, any epoch that contained an auditory frequency distortion, a visual brightening, or a button press were additionally excluded from analyses. In the attention task, we also removed all data from any trial with an error (false alarm or miss). The average number of epochs remaining for each condition is shown in Table 1.

**Table 1.**
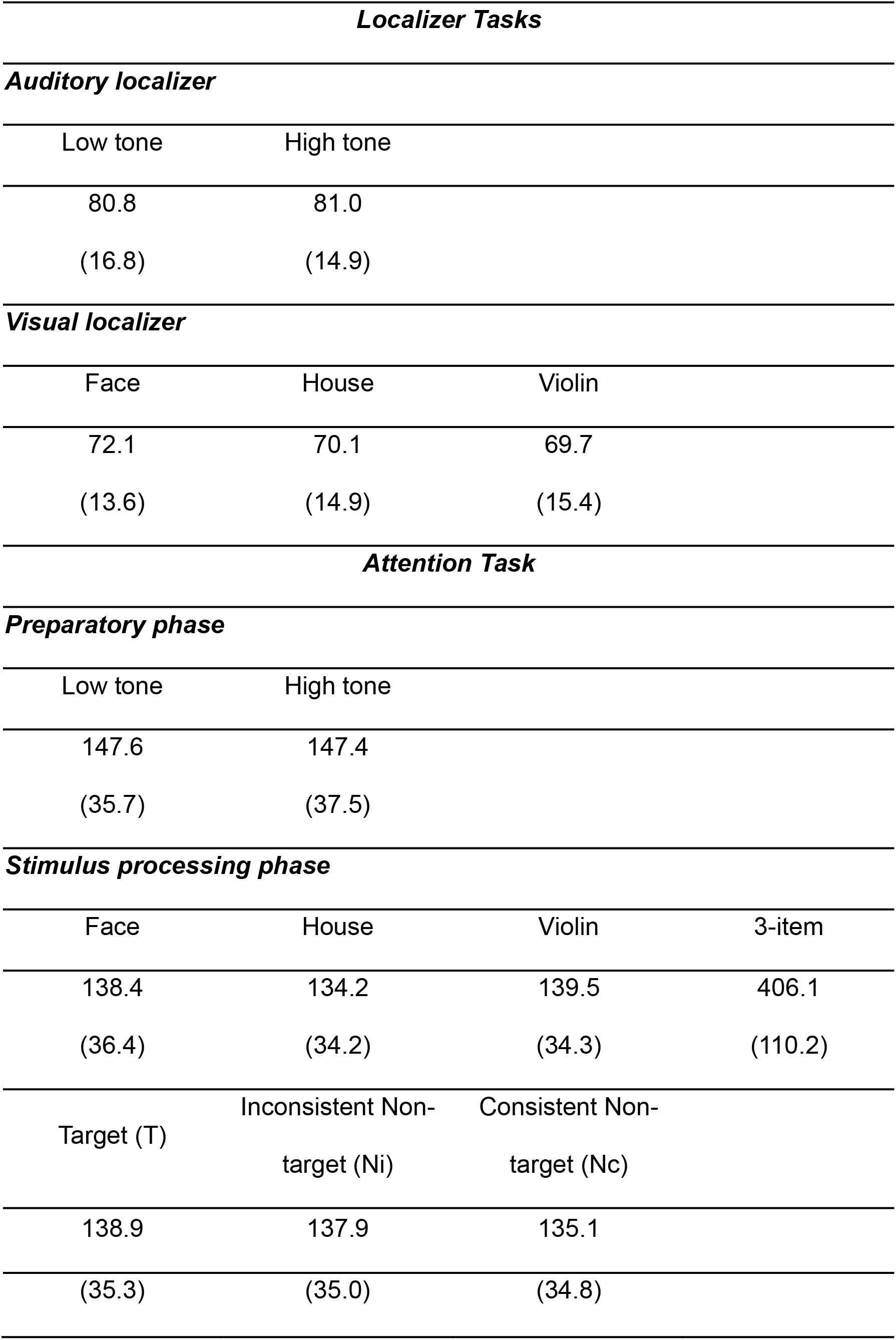
Mean number of epochs (and standard deviation across participants) per condition after artifact rejection

| Localizer Tasks |  |  |  |
| --- | --- | --- | --- |
| Auditory localizer |  |  |  |
| Low tone | High tone |  |  |
| 80.8 | 81.0 |  |  |
| (16.8) | (14.9) |  |  |
| Visual localizer |  |  |  |
| Face | House | Violin |  |
| 72.1 | 70.1 | 69.7 |  |
| (13.6) | (14.9) | (15.4) |  |
| Attention Task |  |  |  |
| Preparatory phase |  |  |  |
| Low tone | High tone |  |  |
| 147.6 | 147.4 |  |  |
| (35.7) | (37.5) |  |  |
| Stimulus processing phase |  |  |  |
| Face | House | Violin | 3-item |
| 138.4 | 134.2 | 139.5 | 406.1 |
| (36.4) | (34.2) | (34.3) | (110.2) |
| Target (T) | Inconsistent Non-target (Ni) | Consistent Non-target (Nc) |  |
| 138.9 | 137.9 | 135.1 |  |

|  |  |  |
| --- | --- | --- |
| (35.3) | (35.0) | (34.8) |

### 2.5 Source Localization

For each participant, a cortical mesh was created from the individual’s structural MRI, with a mesh resolution of ∼4000 vertices per hemisphere. The EEG/MEG and MRI were coregistered based on the three anatomical fiducial points and an additional ∼200 digitized points on the scalp. Forward models were computed for EEG data using a three-shell boundary element model (BEM) and for MEG data using a single-shell BEM. The forward model was inverted using minimum norm estimation (MNE) to compute source estimates for each experimental condition.

Due to the limited spatial resolution limits of EEG/MEG, we chose three *a priori* spatially distinct bilateral ROIs (Figure 2C). Early visual cortex and lateral prefrontal cortex (LPFC) were used to test representation in relevant sensory and cognitive control areas. An additional auditory cortex ROI was used both to measure cue decoding, and in other analyses to test for signal leakage. Auditory and primary visual cortex ROIs were taken from the SPM Anatomy toolbox (Eickhoff et al., 2005), containing 350 and 523 vertices. The LPFC ROI was taken from Fedorenko et al. (2013) (http://imaging.mrc-cbu.cam.ac.uk/imaging/MDsystem), combining the anterior, middle, and posterior medial frontal gyri, spanning 461 vertices.

**Figure 2.**
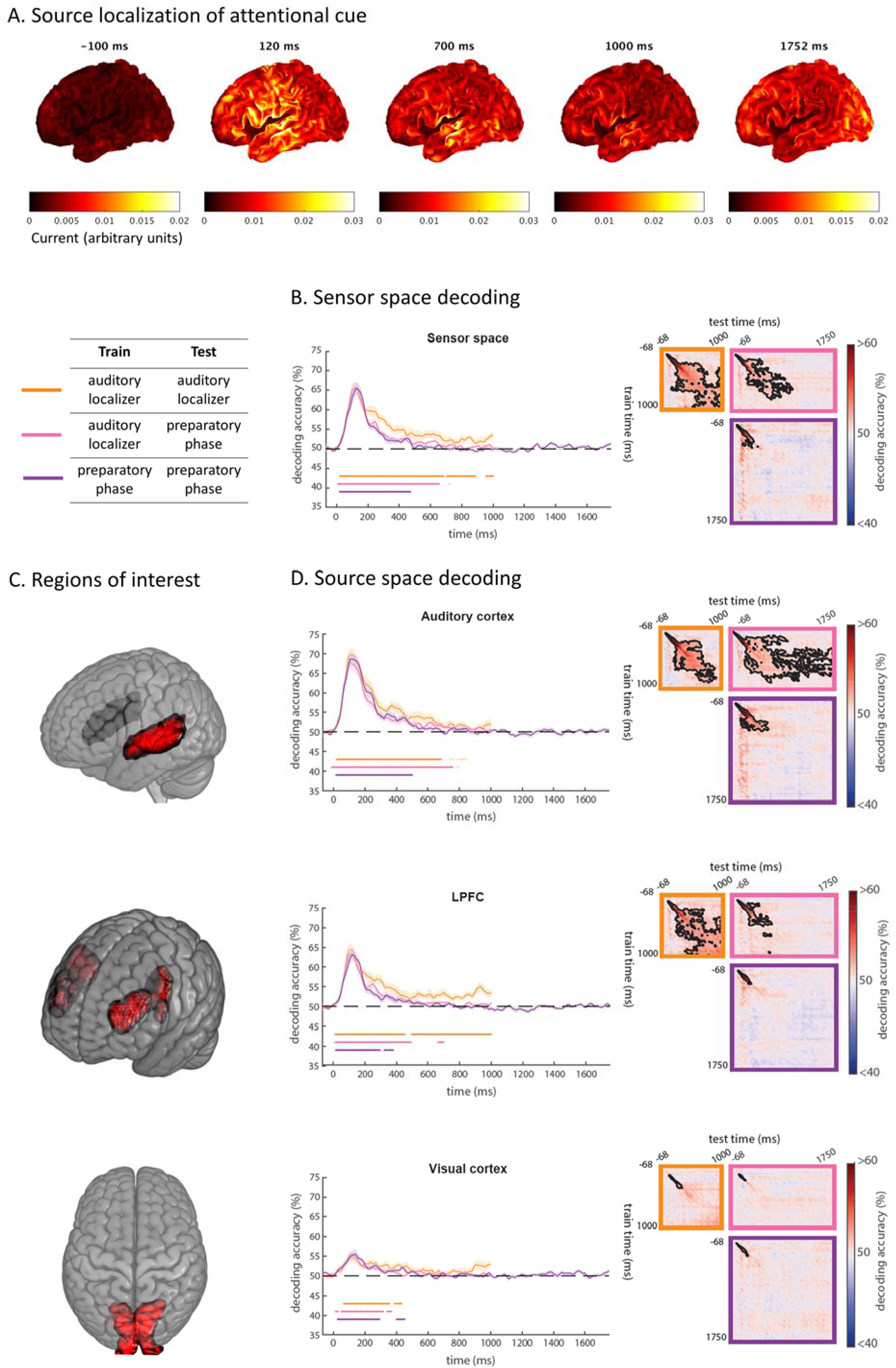
Response to the attentional cue. (A) Source localization of EEG/MEG response to the auditory attentional cue at representative time points relative to cue onset. (B) Decoding time-course of auditory stimulus/attentional cue using all sensors combining EEG and MEG across the whole brain. Curves on the left show decoding when training and testing on matched time-points. Dark colored dots beneath the decoding curves show times where decoding is significantly above chance for each condition (p < 0.05), corrected for multiple comparisons along the diagonal of the cross-temporal generalization matrix; faint colored dots represent additional time-points where the diagonal of the cross-temporal generalization matrix is significant when corrected for multiple comparisons across the whole matrix. Translucent bands represent standard error of the mean. Matrices on the right show temporal generalization of decoding across all pairs of training and testing times. Black contours indicate regions of significant decoding (p<0.05). (C) Vertices within source space ROIs (auditory cortex, lateral prefrontal cortex (LPFC), and visual cortex). (D) Decoding time-courses from these source space ROIs; same format as (B). Significance is corrected for multiple comparisons across time using TFCE and permutation testing.

We chose V1 as the visual ROI to keep the three regions as far apart as possible and thus minimize signal leakage between them. Since higher visual regions are specialized for object-level processing, and can contain template-like signals (Stokes et al., 2009), we subsequently examined a broad extrastriate visual cortex (ESV) ROI from the Fedorenko et al. (2013) template, which encompasses object, face, and scene processing regions. In all cases, results were very similar, reflecting the low spatial resolution of MEG. Here we report the results of V1, but the results from ESV can be found in Supplementary Material 1.

### 2.6 Multivariate Pattern Analysis (MVPA)

Multivariate pattern analyses were performed using the Matlab interface of LIBSVM (Chang and Lin, 2011). We used a linear support vector machine (SVM), with default parameters. For each analysis, we performed decoding in sensor space as well as in source space using data from the three ROIs. For sensor space decoding, we combined data from good EEG and MEG (gradiometers and magnetometers) channels. Each individual time point was standardized (z-scored across channels) before entering the classifier. For source space decoding, each participant’s cortical mesh was transformed into MNI space, and estimated source activity at each vertex within the ROIs was extracted to serve as a feature in the classifier.

In both sensor and source space MVPA analyses, we trained and tested using spatiotemporal patterns extracted from a sliding time window of 32 ms, in 4 ms steps. Training and testing were performed on every combination of time windows, resulting in a cross-temporal generalization matrix of classification accuracies (King and Dehaene, 2014), with the diagonal representing the performance of classifiers trained and tested on the same time window. The classification accuracy matrix was then slightly smoothed using a sliding 32 ms square averaging window. For analyses involving within-task decoding, the data were split into five folds (with one fold containing every 5^th^ trial chronologically), iteratively trained on individual trials from four of the folds and tested on the remaining fold by applying the SVM to the remaining trials individually. In cross-task decoding, a classifier was trained on all relevant epochs from one task and tested on all relevant epochs from another task.

Classification accuracies were compared against chance (50%) with one-tailed t-tests. Multiple comparisons were accounted for using Threshold Free Cluster Enhancement (TFCE), with height and extent exponents of 2 and 2/3 respectively, and Family-Wise Error controlled by comparing the statistic at each time point to the 95^th^ percentile of the maximal statistic across a null distribution of 1000 permutations with random sign flipping (Smith and Nichols, 2009). TFCE was performed in the same way across the time × time decoding matrices and along the matched-time diagonals. The figures were plotted according to the last time bin in the sliding window (Grootswagers et al., 2017). For decoding of 1-item behavioral category, epochs that were preceded by a T or Ni were excluded, to ensure that behavioral category was balanced in the baseline period.

### 2.7 Data and code availability statement

The data and code are available upon direct request of the corresponding author.

## 3. Results

### 3.1 Behavioral results

Behavioral performance was consistently high (auditory localizer task – hits: mean = 99.0%; false alarms: mean = 0.8%; visual localizer task – hits: mean = 98.9%; false alarms: 1.0%; attention task – hits: mean = 98.3%; false alarms: mean = 0.8%).

### 3.2 Coding of the attentional cue/attentional template during the preparatory phase

Source localization of the response to the cue at representative time points is shown in figure 2A. We first looked for decoding of the specific attentional cue during the preparatory phase of the attention task, defined as starting from cue onset but before the first visual stimulus, and compared this with decoding in the auditory localizer task. We asked whether preparing for a target enhances cue decoding. Here, we subsampled the trials in the attention task to match the minimum number of trials in the auditory localizer for each participant, keeping the first n trials, to ensure comparable signal-to-noise ratio across the three decoding analyses. Cue/stimulus decoding as a function of time from auditory stimulus onset is shown in Figure 2B, D. Curves on the left show training and testing on matched time-points. Matrices on the right show generalization of patterns across all pairs of training and testing time windows.

Across the whole sensor space (Figure 2B), significant discrimination between the two auditory stimuli/cues emerged shortly after the presentation of the stimulus, peaking at around 116 ms for the auditory localizer task (Figure 2B, orange curve), 148 ms for the preparatory phase of the attention task (Figure 2B, purple curve), and 112 ms when training the classifier on the localizer task and testing on the attention task (Figure 2B, pink curve). In both sensor space (Figure 2B) and all ROIs (Figure2C-D), cue decoding during the attention task returned to chance level. During the auditory localizer task, cue decoding was more sustained, especially in the LPFC. After matching the number of trials used to train the classifier, an analysis type × ROI ANOVA of peak decoding accuracies, within a 0 – 600 ms time window, showed a main effect of ROI (F(2,34) = 155.2, p < 0.001), but no differences in decoding amplitude (F(2,34) = 3.4, p > 0.5), and no interactions (F(4,68) = 0.7, p = 0.6). Therefore, we found no evidence for template representation beyond the initial auditory representation of the cue.

To test whether activity during any stage of the preparatory phase might reflect the representation of the upcoming trial target, we performed a cross-task and cross-time classification analysis trained using the visual localizer task. At every time window, patterns from the two visual items associated with each cue were taken from the visual localizer task to use as training data, and these were tested at every time window of the preparatory phase of the attention task to decode the trial target (now without subsampling trials). We did not find any significant time points where the visual template cross-generalized to the preparatory phase.

Finally, we note that cross-time generalization matrices suggest that the LPFC signal reached a steady state at the end of the auditory localizer, in contrast to its lack of any sustained signal during the preparatory phase of the attention task. Even including all the trials of the attention task, without subsampling, we observed the same disappearance of cue decoding during the preparatory phase (see Supplementary Material 2). This might reflect that the fact that the representation in the auditory task does not need to be transformed further, whereas in the attention task it serves an intermediate role in mapping subsequent visual inputs to behavior.

### 3.3 Coding of visual and behavioral properties of 1-item displays

We next turned to processing of the visual items, and selection of the target item. Source localization of the response to the visual stimuli at representative time points is shown in figure 3A.

**Figure 3.**
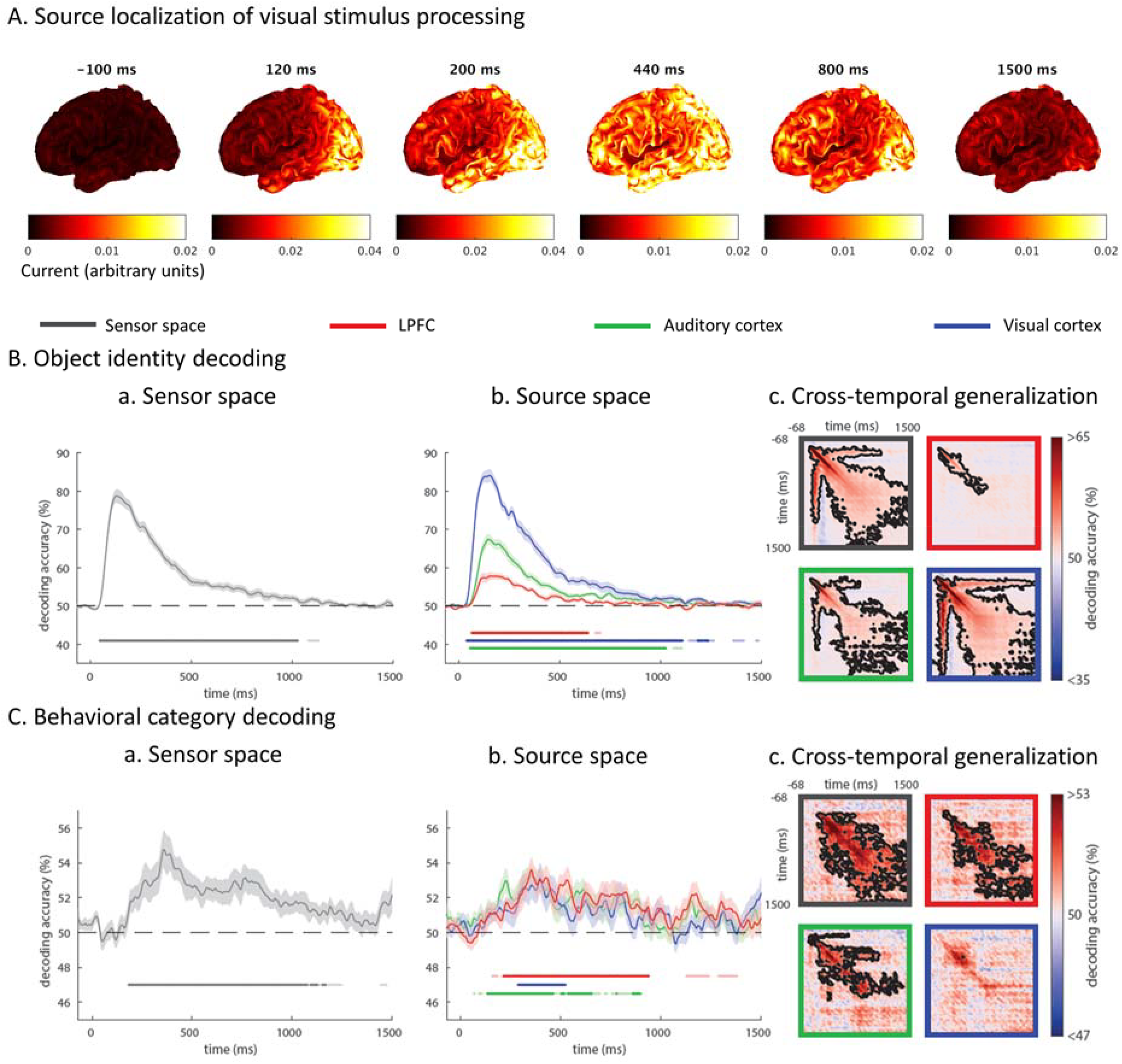
Coding of visual and behavioral properties of 1-item displays (A) Source localization of EEG/MEG response to visual presentation (including both single-item and multi-item displays) at representative time-points. (B) Decoding time-courses of object identity, in (a) sensor and (b) source space, when training/testing using matched time-points, and (c) generalizing across training/testing times. Dark colored dots beneath the decoding curves show times where decoding is significantly above chance for each condition (p < 0.05), corrected for multiple comparisons along the diagonal of the cross-temporal generalization matrix; faint colored dots represent additional time-points where the diagonal of the cross-temporal generalization matrix is significant when corrected for multiple comparisons across the whole matrix. Translucent bands represent standard error of the mean. Black contours in cross-time matrices indicate regions of significant decoding (p<0.05). Significance is corrected for multiple comparisons across time using TFCE and permutation testing. (C) Decoding time-courses and cross-temporal generalization for behavioral category information. Object identity decoding emerged earlier than behavioral category decoding. Visual cortex showed the highest object decoding accuracy, while ROIs were comparable in their strength of behavioral category representation.

During 1-item displays, we expected strong, early discrimination of object identity (e.g., face vs. house, when the consistent non-target was the violin). In the attention task, each stimulus additionally had a behavioral category depending on the cue of that trial. For the participant to make the appropriate response to each stimulus, we expected that the neural signal would also show behavioral category discrimination (target vs. non-target), which would occur after object identity processing. For these analyses, we focused on the T and Ni conditions, for which object identity and behavioral category were fully crossed. Object representation was measured by the discrimination between stimulus identities (e.g. face vs. house) when each were equally often targets or non-targets; conversely, behavioral category representation was measured by discrimination between targets and non-targets when these were equally balanced across stimulus identities.

Single stimulus decoding time-courses on T and Ni presentations are shown in Figure 3B-C. In line with expectations, both object identity and behavioral category showed substantial periods of significant decoding accuracy. Across the whole sensor space, a significant difference between object identities peaked at around 128 ms. Behavioral category decoding emerged later, slowly rising to a peak at 360 ms.

Source space analysis showed that both types of information could be decoded from all three ROIs. Decoding of object identity in the auditory ROI warns of possible signal leakage between regions. Visual cortex, however, had the highest decoding accuracy for object identity, while ROIs did not statistically differ in their strength of decoding accuracy for behavioral category.

Cross-temporal generalization indicated that object identity representation was most stable in the visual ROI. In contrast, behavioral category representation was most stable in the LPFC ROI.

### 3.4 Coding of target location in 3-item displays

Next, we examined target representation in the presence of simultaneous distractors. We first asked when the spatial location of the target within 3-item displays could be decoded (Figure 4; Fahrenfort et al. 2017). To do this, we decoded every pair of T versus Ni locations, while holding Nc position constant (i.e., “T right, Ni left” vs. “Ni right, T left”, “T right, Ni bottom” vs. “Ni right, T bottom”, and “T left, Ni bottom” vs. “Ni left, T bottom”) and averaged the accuracies within each participant. Within each pair, collapsing across both possible cues ensured that the decoding was balanced for both visual features and auditory cues. Group sensor-space results showed that decoding began to emerge shortly after stimulus onset, and peaked at 244 ms, before slowly declining toward the end of the epoch. The analysis was repeated in source space. Decoding of target location was significant in all ROIs, but strongest in visual cortex where it peaked at 132 ms. Cross-temporal generalization suggested that the representation of target location was initially dynamic, then entered a temporarily stable state, most apparent in sensor space suggesting spatially coarse stability, before becoming unstable once more prior to the end of the epoch.

**Figure 4.**
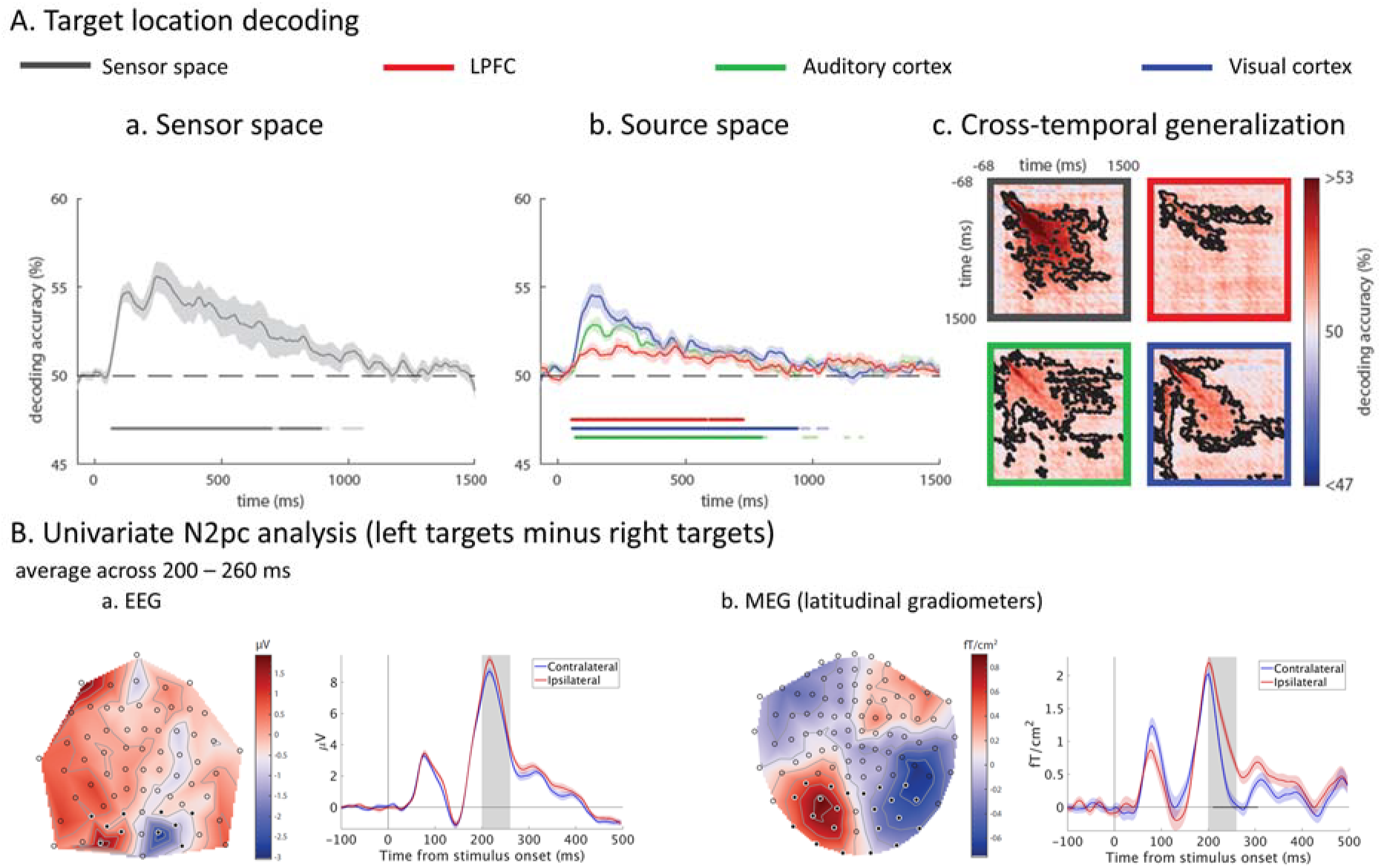
Coding of target location in 3-item displays. (A) Decoding of target location during presentation of 3-item displays, i.e., whether the item corresponding to the cue is in the left, right, or bottom position. Format as in Figure 3. Location decoding was strongest in the visual cortex. (B) Univariate N2pc ERP/ERFs across (a) EEG electrodes and (b) latitudinal gradiometers. Latitudinal gradiometers are presented because their orientation around the helmet means that contralateral asymmetries in the magnetic flux gradient are expressed analogously to the EEG topography (Mitchell and Cusack, 2011; Kuo et al. 2016). Topographies are averaged across 200-260 ms (marked in grey on the time-courses). Time-courses are averaged across posterior sensors contralateral and ipsilateral to the target (highlighted on the topographies), with black dots indicating a significant difference (p<0.05) after TFCE with permutation testing.

In a complementary analysis to target location decoding, we examined the N2pc, a well-known early index of spatial attention, which appears as a negativity over posterior EEG electrodes contralateral to the side of space to which the subject is attending around 200-300 ms following a stimulus (Luck and Hillyard, 1994; Heinze et al., 1990; Eimer, 1996; Hopf et al. 2000; Fahrenfort et al., 2017). We compared event-related potentials/fields when the target was on the right or left of the screen of the 3-item display, and the topography of this contrast is shown in Figure 4B. Differences between target locations peaked between 200-300 ms in posterior EEG and MEG signals, although the signals diverged earlier in MEG, which could reflect the source of the earlier decoding. We note that our lateralized stimuli were in the upper visual field, and that the N2pc is typically stronger for stimuli in the lower visual field (Luck et al 1997; Bacigalupo & Luck, 2018).

### 3.5 Coding of target identity during presentation of 3-item displays

We also hypothesized that representation of 3-item displays would differ depending on the cue, even though the visual input was the same. All 3-item displays contained the target item that was associated with the cue, as well as the Ni and Nc items. Therefore, the decoding of the cue in the presence of a matching visual stimulus likely reflects attentional enhancement of the selected target identity. Although a template representation could also contribute to the decoding, this can only be isolated in the absence of a target (see next section). In sensor space, cue/target identity decoding peaked at 252 ms. In source space, the visual cortex showed the highest decoding accuracy (Figure 5).

**Figure 5.**
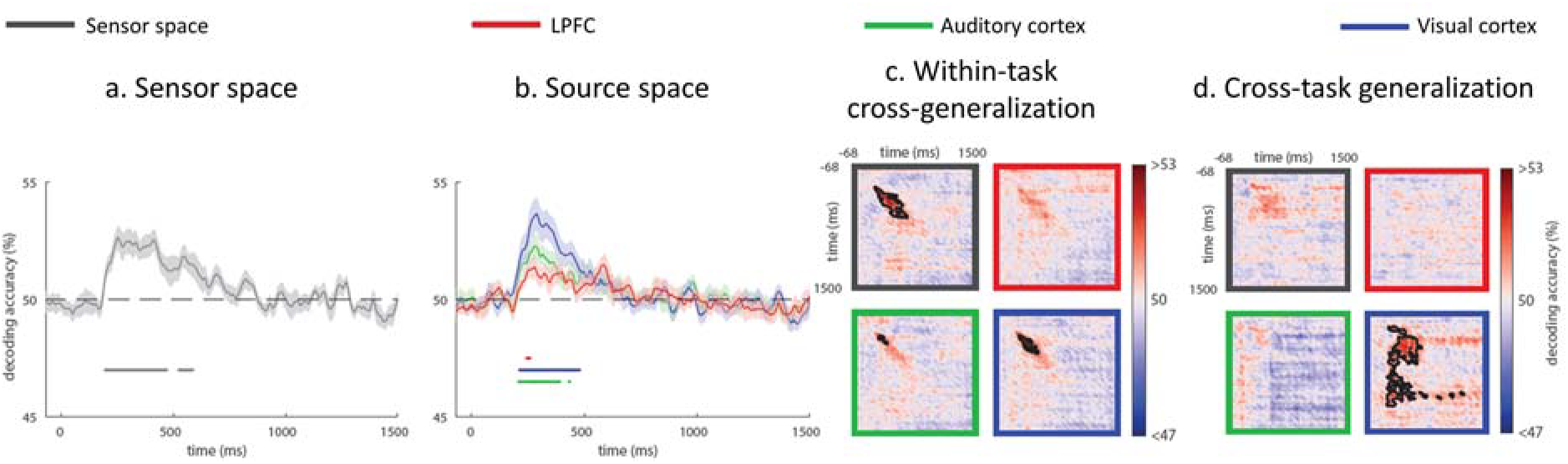
Decoding of attentional cue/target identity during presentation of 3-item displays. Panels (a-c) have the same format as Figure 3. Panel (d) shows cross-task generalization of decoding, when training on the visual localizer task and testing on the attention task.

Cross-temporal generalization suggested that the representation of target identity in the presence of distractors was dynamic, and decayed rather than settling into a steady state. For this analysis, we also expected cross-task generalization from the visual localizer. This was significant in the visual ROI, but not in the auditory or LPFC ROIs, suggesting that the shared pattern was predominantly sensory, with minimal signal leakage in this case.

To compare the decoding latencies of target location and target identity in 3-item displays, we calculated 50%-area latency (Luck, 2014; Liesefeld, 2018) using data from a 0 – 600 ms window for each subject, ROI and decoding type. Paired 2-tailed t-tests showed that target location decoding preceded target identity decoding in both the whole sensor space (t(17) = 2.86, p < 0.05) and in the visual cortex (t(17) = 4.97, p < 0.001), but not in the auditory or LPFC ROIs (both t(17)<1.95; both p>0.05).

### 3.6 Reawakening of the attentional cue/template during presentation of consistent non-targets

Finally, we tested whether we could decode the cue/template during the presentation of a single Nc visual stimulus. Wolff et al. (2015, 2017) have shown that by ‘pinging’ the brain with a neutral stimulus during working memory maintenance, it can be possible to decode the memory-item-specific information from the impulse response. In our data, cue decoding following Nc presentation was visible but rather weak and intermittent (Figure 6). Across sensor space and source space, there were scattered brief periods of above-chance decoding. Their appearance in auditory as well as visual and frontal ROIs questions whether these might reflect a reactivated memory of the auditory cue, or a visual attentional template in anticipation of the next visual input. Apparent signal in the auditory ROI might also reflect leakage from other sources. Cross-temporal generalization suggested that although the representation was not fully sustained, when it resurfaced in the visual ROI it did so with a similar pattern. Cross-task generalization from the auditory and visual localizers provided no evidence that this representation was in a similar format to either cue or target perception.

**Figure 6.**
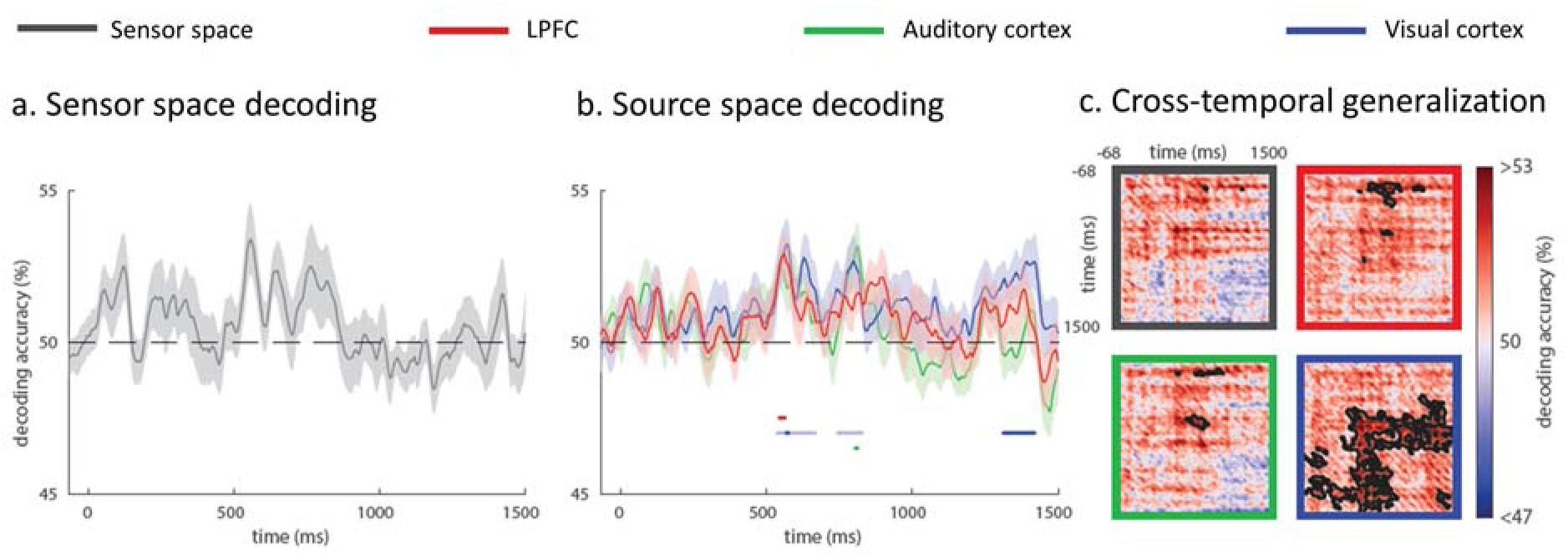
Decoding time-course of attentional cue during presentation of Nc displays. Format as in Figure 3.

### 3.7 Summary of component time-courses during attentional selection

Above we have described five distinct forms of information representation evoked by the appearance of the visual stimuli (Figures 3–6). These are summarized in Figure 7, overlaying their average sensor-space and ROI-based decoding time-courses for ease of comparison.

**Figure 7.**
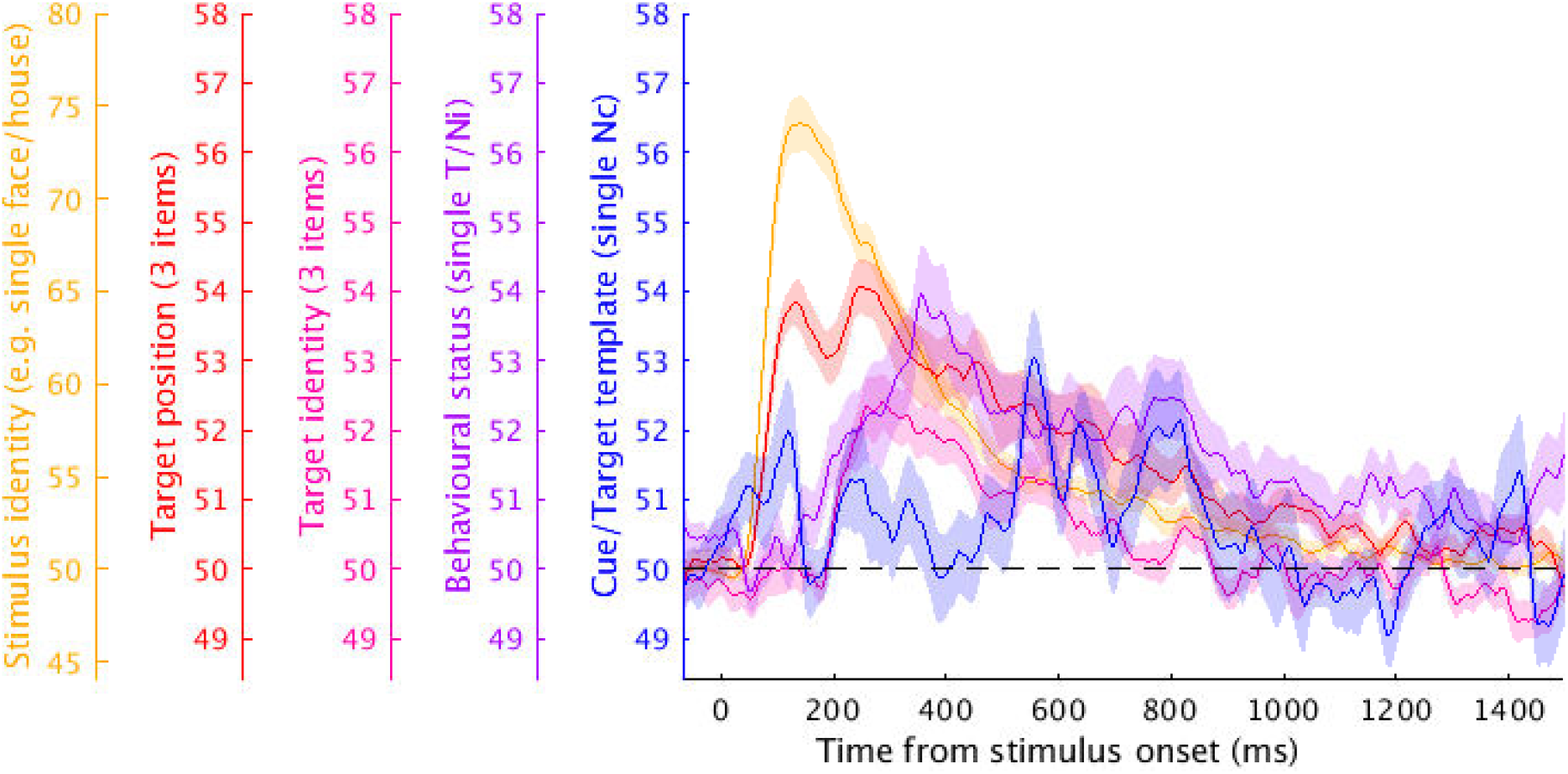
Summary of the decoding time-courses of five component processes of selective attention following onset of a visual stimulus: representation of stimulus identity, target position, target identity, behavioral status, and the template of the cue/target. Decoding accuracy is averaged across sensor space and source ROIs, and translucent bands represent standard error of the mean across subjects.

## 4. Discussion

There is currently much interest in decoding the contents of cognitive operations from human MEG/EEG data, and in using these methods to understand attentional selection of information relevant to current goals. Here, we examined the evolution of multiple forms of information represented in the brain as a visual target is selected. Combining single-item displays with multi-item displays of targets and different types of distractors allowed quantification of distinct components of processing during selective attention, indexed by different profiles of representational content.

Although multiple attentional templates could guide behavior (Awh et al., 2012), for effective task performance selection of a particular target requires a template that specifies the currently relevant object (Duncan and Humphreys, 1989; Bundesen, 1990). In fMRI, multivariate classifiers trained on responses to viewed stimuli can predict an attentional template during the preparatory phase (e.g., Stokes et al., 2009). In our MEG/EEG data, we observed significant decoding of cue identity in the attention task, but after equating trial numbers decoding accuracies were not significantly different from that of stimulus processing in the auditory localizer task. Furthermore, beyond 1000 ms, cue decoding was indistinguishable from chance. Previous MEG/EEG studies have suggested existence of a pre-stimulus template, often subtle and short-lived (Myers et al., 2015; Kok et al., 2017; Grubert and Eimer, 2018). Following non-target displays, we observed evidence of template reawakening; although significant, this was weak and not fully sustained. The delay of template reactivation relative to the explicit categorization of the display as a non-target suggests a serial component to the search process, here within the temporal presentation stream but consistent with neural evidence of serial refocusing of attention within single search displays (Woodman & Luck, 1999; Bichot et al., 2005).

Sustained preparatory activity reflecting an attentional template may be largely invisible to MEG/EEG for many reasons. For example, at the physiological level, if discriminating neurons are intermixed, they may be hard to distinguish with non-invasive methods. Recent findings from single trial analysis of direct neural recordings also suggest that spiking activity during the delay period is sparse, with brief bursts of activity having variable onset latency and duration, which would hinder cross-trial decoding (Shafi et al., 2007; Lundqvist et al., 2016, 2018; Stokes and Spaak, 2016; Miller et al., 2018). A parallel possibility is that attentional templates may sometimes be stored in an “activity silent” passive form, such as changed synaptic weights (Lundqvist et al., 2010; Stokes, 2015). Consistent template representations may also be difficult to detect if there is-trial-to-trial variability at the cognitive level (Vidaurre et al., 2019), such as fidelity of mental imagery, as well as the anticipation of stimulus timing, with templates activated/strengthened only when the search display is expected to be imminent (Grubert and Eimer, 2018). It is also possible that in the current experiment, the attentional template required little effort to maintain as a verbal label and might have been more visible if harder to verbalize. Consistent templates may be more likely when few features distinguish targets from distractors, for example when targets are defined only by orientation or color (Kok et al., 2017; Myers et al., 2015; Grubert and Eimer, 2018). Perceptually complete templates may be more likely when targets share different features with different distractors (Duncan and Humphreys, 1989). Finally, we emphasize that for successful task performance a template must exist in some form, even when we are unable to detect it, and that uncovering subtle or variable templates may benefit from novel analysis methods (Vidaurre et al. 2019).

Upon presentation of the visual choice display, we found much decodable information of various kinds. The timing of peak decoding of different features suggests five components of processing. The current data cannot determine the extent to which these components evolve in parallel or have some serial dependency, whereby one process influences another. It is likely that there is a degree of both (Bichot et al., 2005). First, visual stimulus properties are encoded, shown by object identity decoding in 1-item displays, peaking around 132 ms, and strongest in visual cortex. Second, in a multi-element display, the candidate target is localized, shown by target location decoding that peaked between 136 ms (in visual cortex, where strongest) and 288 ms (combining all sensors). This may be partially concurrent with initial visual processing, consistent with an initial parallel stage of selection (Duncan, 1980; Treisman and Gelade, 1980) and automatic registration of coarse feature location (e.g. Cohen & Ivry,1989; Hopf et al., 2004), that could be used to guide subsequent attention (Itti and Koch, 2000; Bisley and Goldberg, 2010; Wolfe, 1994; Eimer, 2015). Third, representation of the candidate target continues to be enhanced relative to distractors, perhaps via integrated competition, shown by cue/target identity decoding in 3-item displays, peaking around 252 ms, again strongest in visual cortex. Fourth, behavioral significance of the target is explicitly represented (in this case whether it is a target, so requiring further processing), shown by behavioral category decoding in 1-item displays, peaking around 344 ms and most stable in the LPFC. Fifth, if no target is identified and search must continue, an attentional template might be reactivated or strengthened, shown by cue decoding after Nc displays, peaking beyond 500 ms. The precise timing at which each representation is detectable will depend on many factors including stimuli, task, analysis sensitivity, similarity between targets and distractors, and the number and homogeneity of distractors (Duncan and Humphreys 1989). Nonetheless, we anticipate that the sequence of key components would largely generalize across paradigms (Eimer, 2015; Vidaurre et al., 2019). Potential dependencies between processes might be investigated by combining MVPA of electrophysiological recordings with transcranial magnetic stimulation at successive times.

In 1-item displays, we found a distinction between visual cortex and LPFC. While the regions represented behavioral category with similar strength, visual cortex represented stimulus identity more strongly than LPFC. Similarly, object identity was represented more stably in visual cortex, whereas behavioral category was represented more stably in LPFC. fMRI studies show that frontal regions flexibly code for behaviorally relevant categories according to task rule (Jiang et al., 2007; Li et al., 2007; Woolgar et al., 2011; Lee et al., 2013; Erez and Duncan, 2015). Electrophysiological recordings of monkey prefrontal responses to T, Ni, and Nc stimuli show that visual input properties are initially equally represented for targets and non-targets, whereas the behaviorally critical target dominates later processing (Kadohisa et al., 2013; Stokes et al., 2013). Our results also suggest an anterior-posterior distinction in information content and timing.

In multi-item displays, the candidate target was rapidly identified and localized, with location decoding providing the earliest evidence of modulation by behavioral relevance. Although it’s timing, peaking around 132 ms in V1, was earlier than might be expected based on the N2pc and multivariate decoding using EEG alone (Fahrenfort et al., 2017), it is consistent with representation of the location of task-relevant features reported from ∼140 ms and preceding the N2pc (Hopf et al. 2004). Ipsilateral and contralateral target responses diverged earlier in MEG than EEG, suggesting that the source of the earlier decoding may be more visible to MEG. Location decoding peaked later in the other ROIs and at the sensor level (beyond 230 ms) suggesting that source localization may have helped in isolating the earlier signal.

Although target localization implies target identification, and time-courses of location and identity representation in 3-item displays were heavily overlapping, the location signal was significantly earlier than the identity representation in visual cortex. This is consistent with models of visual attention as well as empirical data that make an explicit distinction between feature selection, where attention is rapidly allocated to candidate objects (Broadbent, 1958), and object recognition, which takes place at a subsequent stage where the features of objects are integrated and their identity becomes accessible (Eimer, 2015; Eimer & Grubert, 2014; Kiss et al., 2013). It could also arise within a continuous competitive framework, without explicit recognition, if neurons representing identity have overlapping receptive fields such that competition amongst them is slower to resolve or benefits from prior spatial filtering (Luck et al. 1997); or if complete identity representation involves several features whose integration is strongly mediated by shared location within spatiotopic maps (Treisman & Zhang, 2006; Schneegans and Bays, 2017). The location of an attended feature can also be represented before the location of a target itself (Hopf et al., 2004), and the temporal priority with which different features of the target are enhanced may depend on the cortical location as well as the particular task demands (Hopf et al., 2005). The observations that competitive representations of target location and target identity peaked at different times, and that neither appeared to reach a permanent steady state, together indicate that the early phase of integrated competition is dynamic, with different aspects of the target representation waxing and waning at different times. In contrast, the later explicit representation of target status settled into a steady state in LPFC that persisted until the end of the epoch.

Interestingly, a target influenced bias in the 3-item displays well before its target status was explicitly decodable in the single-item displays. This strongly suggests at least two stages of target processing, consistent with behavioral manipulations suggesting that spatial selection and target identification are separable (Ghorashi et al., 2010). Distinction between an early, parallel processing stage and a later capacity-limited stage is central to most models of attention (Duncan, 1980; Treisman & Gelade, 1980). Target decoding in 3-item displays peaked at 252 ms with first significance at 196 ms, similar to attentional modulation of stimulus category processing in cluttered scenes observed from 180 ms (Kaiser et al., 2016), and to demonstration of feature-binding during integrated competition (Schoenfeld et al. 2003). The later stage indexed by single-item decoding may correspond to capacity-limited individuation of the integrated target object, allowing its bound properties to become accessible for further processing and goal-directed action (Duncan, 1980; Bichot et al., 2005; Mitchell and Cusack 2008; Christie et al., 2015), in this case likely including the brightness judgement. These two stages could also be interpreted in terms of the “global neuronal workspace” model - the earlier attentional bias reflecting accumulation of pre-conscious sensory evidence; the later explicit representation of target status reflecting conscious awareness and “ignition” of fronto-parietal networks, linked to P3 waves around 300-500 ms (Dehaene & Changuex, 2011; Sergent et al., 2005) and consistent with the timing of peak decoding at 360 ms.

To conclude, although attentional selection must begin with a template, this may be weakly or variably represented (Duncan et al., 1997; Lundqvist et al., 2018; Miller et al., 2018), such that it is largely invisible to MEG/EEG, or even maintained in “silent” form (Stokes, 2015). In agreement with others (Olivers et al., 2011; Myers et al., 2015; Grubert and Eimer, 2018), we suggest that the template may be actively and consistently represented only when needed, and least likely to interfere with other concurrent processes. Integrated competition accounts of attention imply that the template need be neither complete nor constant across trials, consistent with no significant response pattern generalization between template representations and the visual localizer. In contrast, integrated competition suggests that attentional selection and enhancement of stimulus representations will be strong and widespread. Supporting such models, we observed robust, time-resolved decoding of the critical processing stages required to select and enhance a target amongst competing distractors, and to categorize it according to behavioral requirements.

## 5. Acknowledgements

The authors would like to thank Seyedeh-Rezvan Farahibozorg, Tim Kietzmann, Olaf Hauk, Peter Watson, Courtney Spoerer, Melek Karadag, and Darren Price for helpful discussions. This work was supported by funding from the Medical Research Council (United Kingdom), program SUAG/002/RG91365. TW was supported by the Taiwan Cambridge Scholarship from the Cambridge Commonwealth, European & International Trust and the Percy Lander studentship from Downing College.

