## Supplementary Material 1 for "The time-course of component processes of selective attention"

Since complex objects were used in the experiment, and higher visual regions are specialized for object-level processing, we additionally examined a broad extrastriate visual cortex (ESV) ROI from the Fedorenko et al. (2013) template, which encompasses object, face, and scene processing regions. In all cases, results were very similar to those from V1, presumably reflecting the relatively low resolution of EEG/MEG. We report the results from ESV here.

***Coding of the attentional cue/attentional template during the preparatory phase***

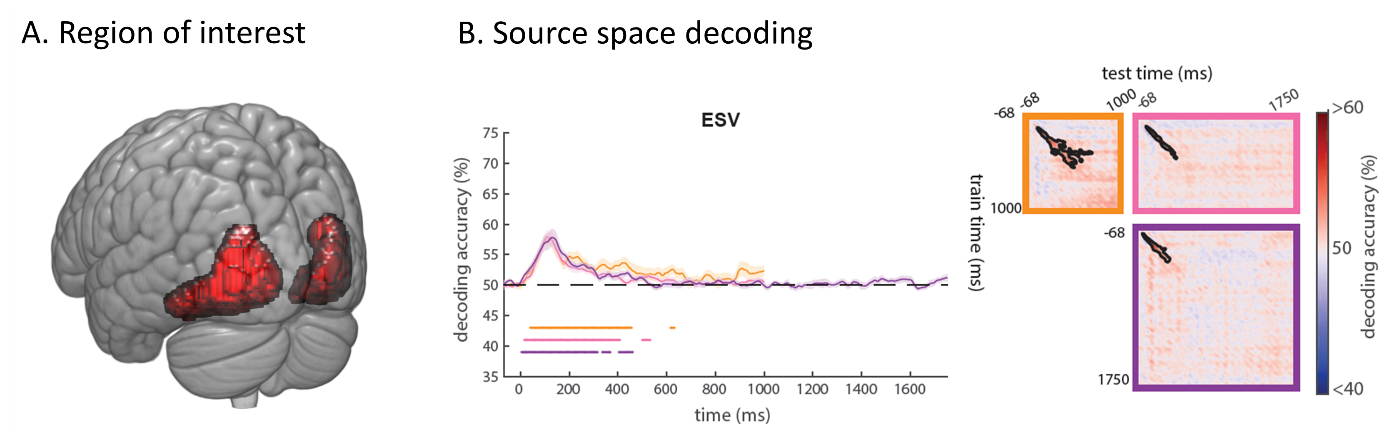

*Supplementary Figure 1. (A) Vertices within source space ESV ROI. (B) Decoding time-course of auditory stimulus/attentional cue from ESV ROI. Curves on the left show decoding when training and testing on matched time-points. Colored dots beneath the decoding curves show times where decoding on the diagonal is significantly above chance for each condition (p < 0.05). Translucent bands represent standard error of the mean. Matrices on the right show temporal generalization of decoding across all pairs of training and testing times. Black contours indicate regions of significant decoding (p<0.05). Significance is corrected for multiple comparisons* across time using TFCE and permutation testing.

To test whether activity during any stage of the preparatory phase might reflect the representation of the upcoming trial target, we performed a cross-task and cross-time classification analysis trained using the visual localizer task. Replicating the results of the three ROIs used in the main experiment, we did not find any significant clusters where the visual template cross-generalized to the preparatory phase in ESV.

***Coding of visual properties of 1-item displays***

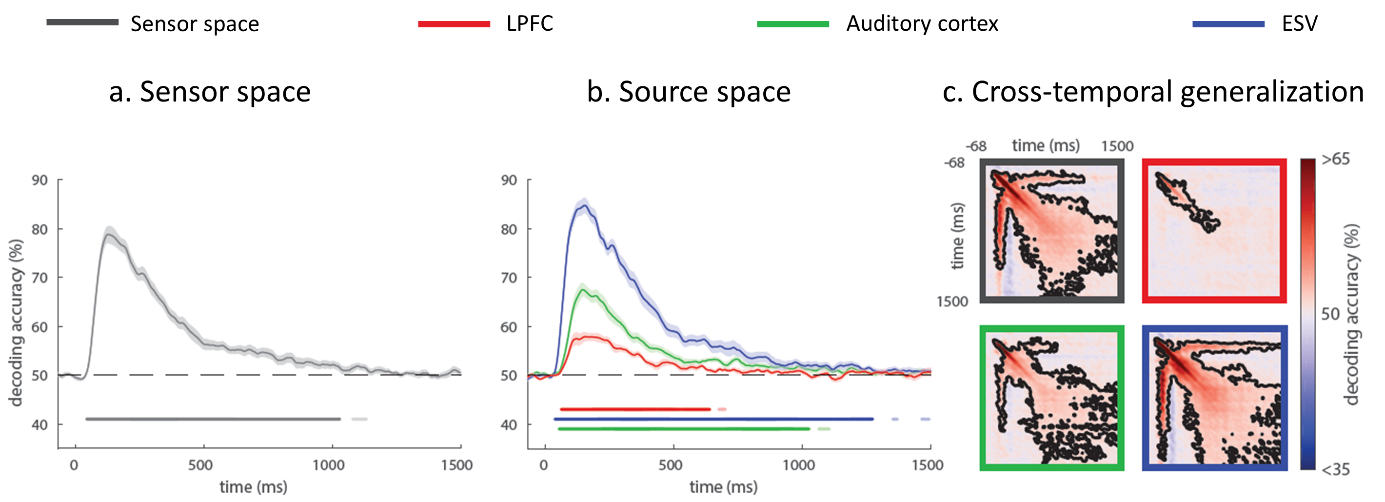

*Supplementary Figure 2. Coding of visual properties of 1-item displays. Decoding time-courses of object identity, in (a) sensor and (b) source space, when training/testing using matched time-points, and (c) generalizing across training/testing times. Dark colored dots beneath the decoding curves show times where decoding is significantly above chance for each condition (p < 0.05), corrected for multiple comparisons along the diagonal of the cross-temporal generalization matrix; faint colored dots represent additional time-points where the diagonal of the cross-temporal generalization matrix is significant when corrected for multiple comparisons across the whole matrix. Translucent bands represent standard error of the mean. Matrices on the right show temporal generalization of decoding across all pairs of training and testing times. Black contours indicate regions of significant decoding (p<0.05). Significance is corrected for multiple comparisons across time using TFCE and permutation testing.*

***Coding of behavioral properties of 1-item displays***

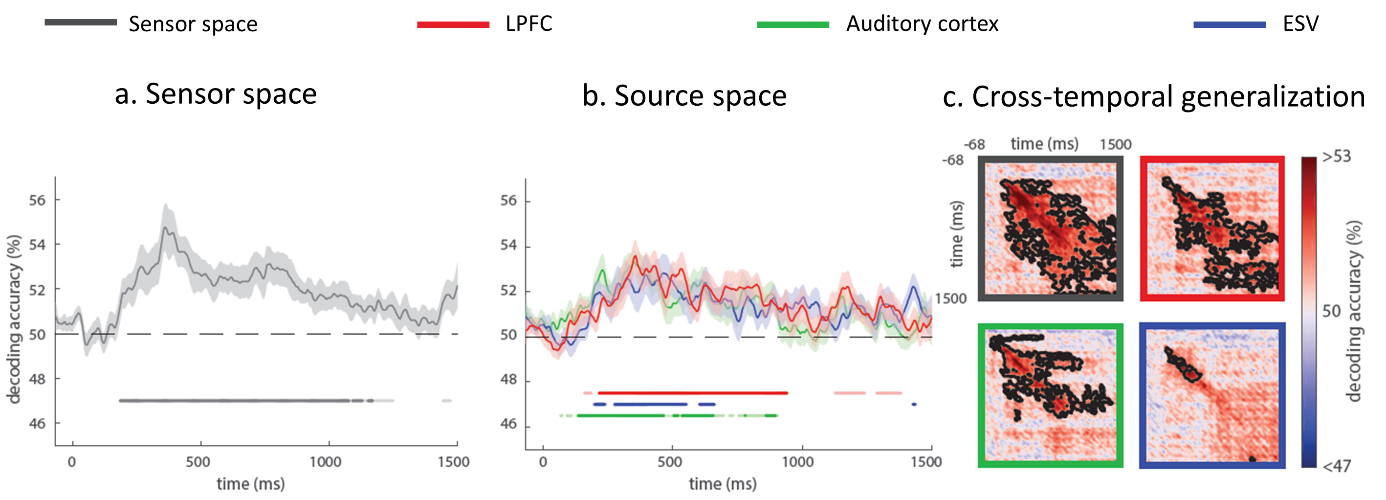

*Supplementary Figure 3. Coding of behavioral properties of 1-item displays. Same format as figure above.*

***Coding of target location in 3-item displays***

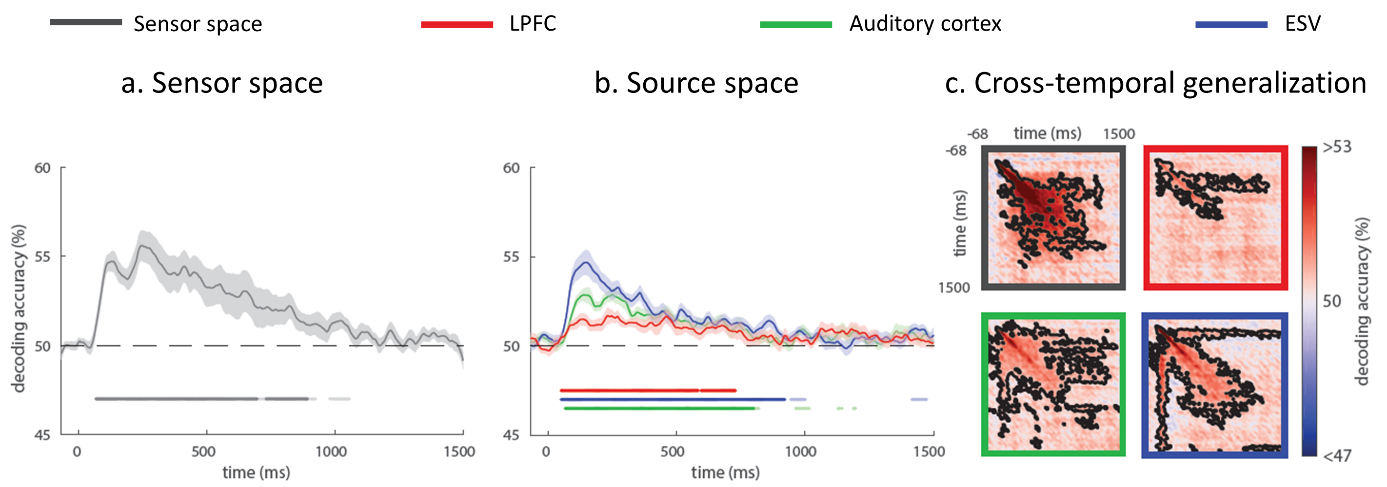

*Supplementary Figure 4. Coding of target location in 3-item displays. Same format as figure above.*

***Coding of target identity during presentation of 3-item displays***

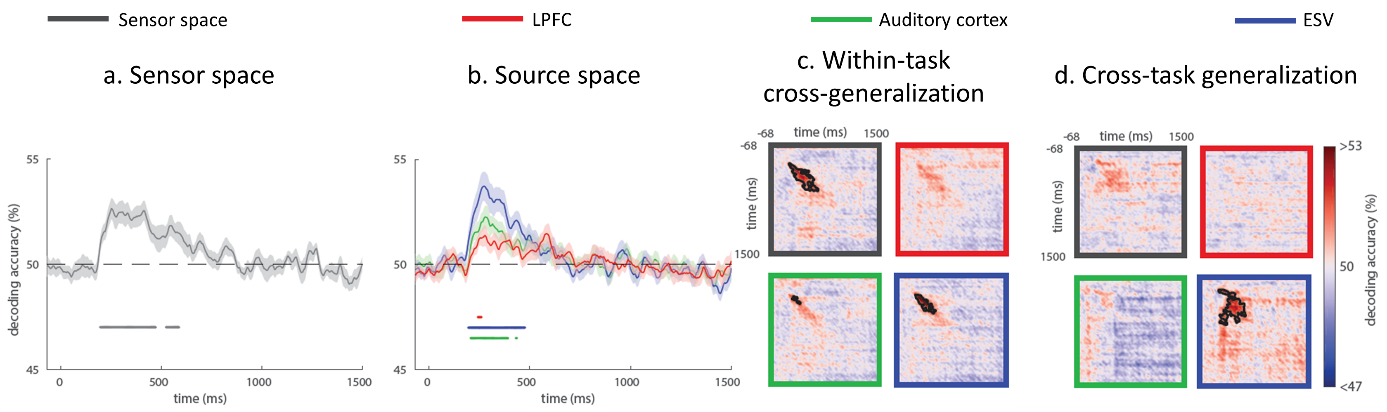

*Supplementary Figure 5.* *Decoding of attentional cue/target identity during presentation of 3-item displays. Same format as figure above.*

***Reawakening of the attentional cue/template during presentation of consistent non-targets***

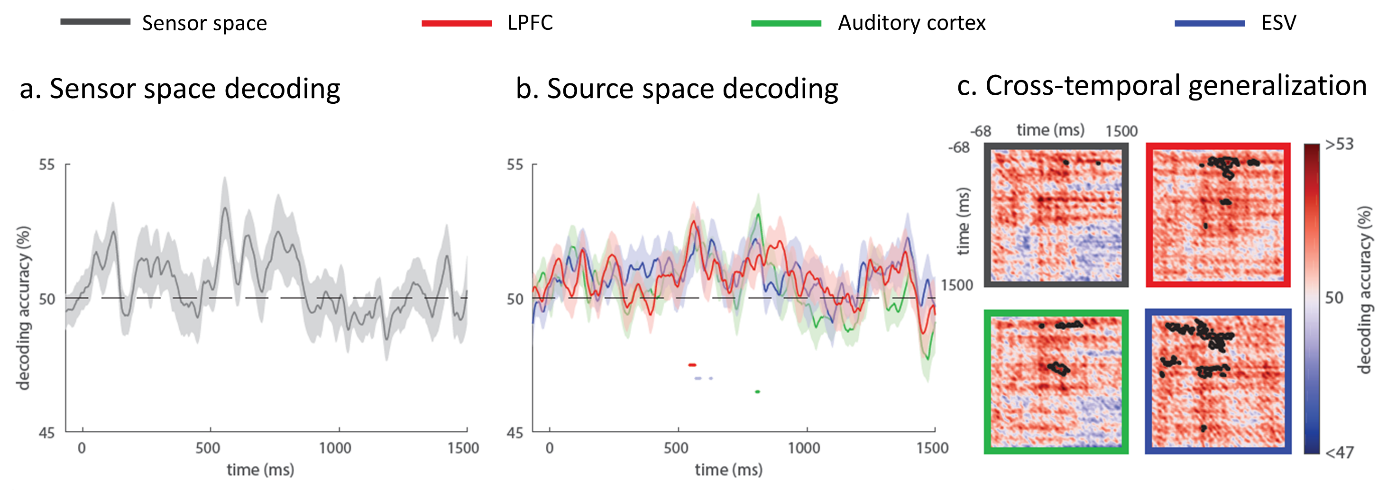

*Supplementary Figure 6.* *Decoding time-course of attention cue during presentation of Nc displays. Same format as figure above.*
