## Supplementary Material 2 for "The time-course of component processes of selective attention"

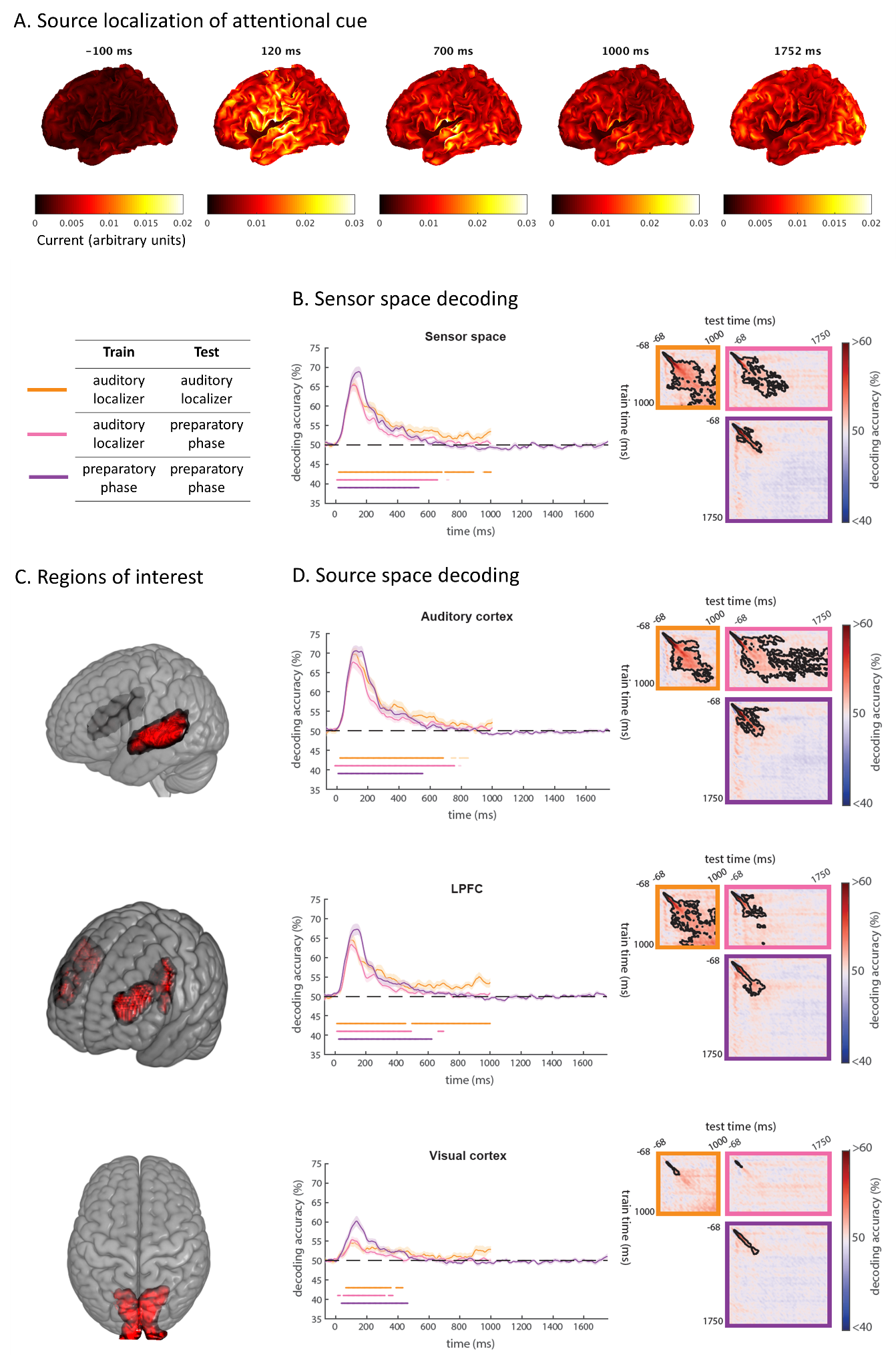

*Supplementary Figure 7. Response to the attentional cue without subsampling the preparatory phase data. (A) Source localization of EEG/MEG response to the auditory attentional cue at representative time points relative to cue onset. (B) Decoding time-course of auditory stimulus/attentional cue using all sensors combining EEG and MEG across the whole brain. Curves on the left show decoding when training and testing on matched time-points. Dark colored dots beneath the decoding curves show times where decoding is significantly above chance for each condition (p < 0.05), corrected for multiple comparisons along the diagonal of the cross-temporal generalization matrix; faint colored dots represent additional time-points where the diagonal of the cross-temporal generalization matrix is significant when corrected for multiple comparisons across the whole matrix Translucent bands represent standard error of the mean. Matrices on the right show temporal generalization of decoding across all pairs of training and testing times. Black contours indicate regions of significant decoding (p<0.05). (C) Vertices within source space ROIs (auditory cortex, lateral prefrontal cortex (LPFC), and visual cortex). (D) Decoding time-courses from these source space ROIs; same format as (B). Significance is corrected for multiple comparisons across time using TFCE and permutation testing.*
